# Taxon cycles in Neotropical mangroves

**DOI:** 10.1101/2022.09.25.509361

**Authors:** Valentí Rull

## Abstract

The concept of taxon cycle involves successive range expansions and contractions over time through which a species can indefinitely maintain its core distribution. Otherwise, it becomes extinct. Taxon cycles have been defined mostly for tropical island faunas, examples from continental areas are scarce and similar case studies for plants remain unknown. Most taxon cycles have been identified on the basis of phylogeographic studies, and straightforward empirical evidence from fossils is lacking. Here, empirical fossil evidence is provided for recurrent Eocene to present expansion/contraction cycles in a mangrove taxon (*Pelliciera*), after a Neotropical-wide study of the available pollen records. This recurrent behavior is compatible with the concept of taxon cycle from biogeographical, chronological and ecological perspectives. The biotic and abiotic drivers potentially involved in the initiation and maintenance of the *Pelliciera* expansion/contraction cycles are analyzed, and the ecological and evolutionary implications are discussed. Whether this could be a trend toward extinction is considered under the predictions of the taxon cycle theory. The recurrent expansion and contraction cycles identified for *Pelliciera* have strong potential for being the first empirically and unequivocally documented taxon cycles and likely the only taxon cycles documented to date for plants.

## 1. Introduction

The concept of taxon cycle was introduced by Wilson (1961) to describe the biogeographical and evolutionary dynamics of species that experience successive range expansions and contractions over time linked to adaptive ecological shifts. According to this author, a taxon can maintain its core distribution area, which he called the headquarters, in a given land mass indefinitely by expanding and contracting its geographical range recurrently. Otherwise, it becomes extinct. In the taxon cycle, expanding and contracting species populations have disparate geographical patterns and adaptive features that allow subdivision of the process into four main stages (Ricklefs & Cox, 1972) (Fig. 1A). In stage I, high-density expanding populations rapidly colonize new environments but bear low morphological differentiation across their geographical range. These taxa have high reproductive potential and broad habitat tolerance, and have been called supertramps (Diamond, 1974). Expansion slows down in stage II, and population differentiation significantly increases, especially near the range margins. The taxa corresponding to this stage are known as great speciators (Diamond et al., 1976). Stage III is characterized by geographical stasis and local extinction leading to fragmented distributions and incipient speciation, which may trigger the onset of a new cycle. If this is not the case, a gradual decline in range size and intraspecific diversity takes place, leading to a progressive relictualization (stage IV) and eventually to extinction (Pepke et al., 2019). According to Ricklefs & Bermingham (2002), the main contribution of the taxon cycle concept to biogeography is the focus on the evolutionary consequences of ecological interactions among colonizing and autochthonous (resident) species, which influence their extinction dynamics and shape their geographical distribution patterns.

**Figure 1.**
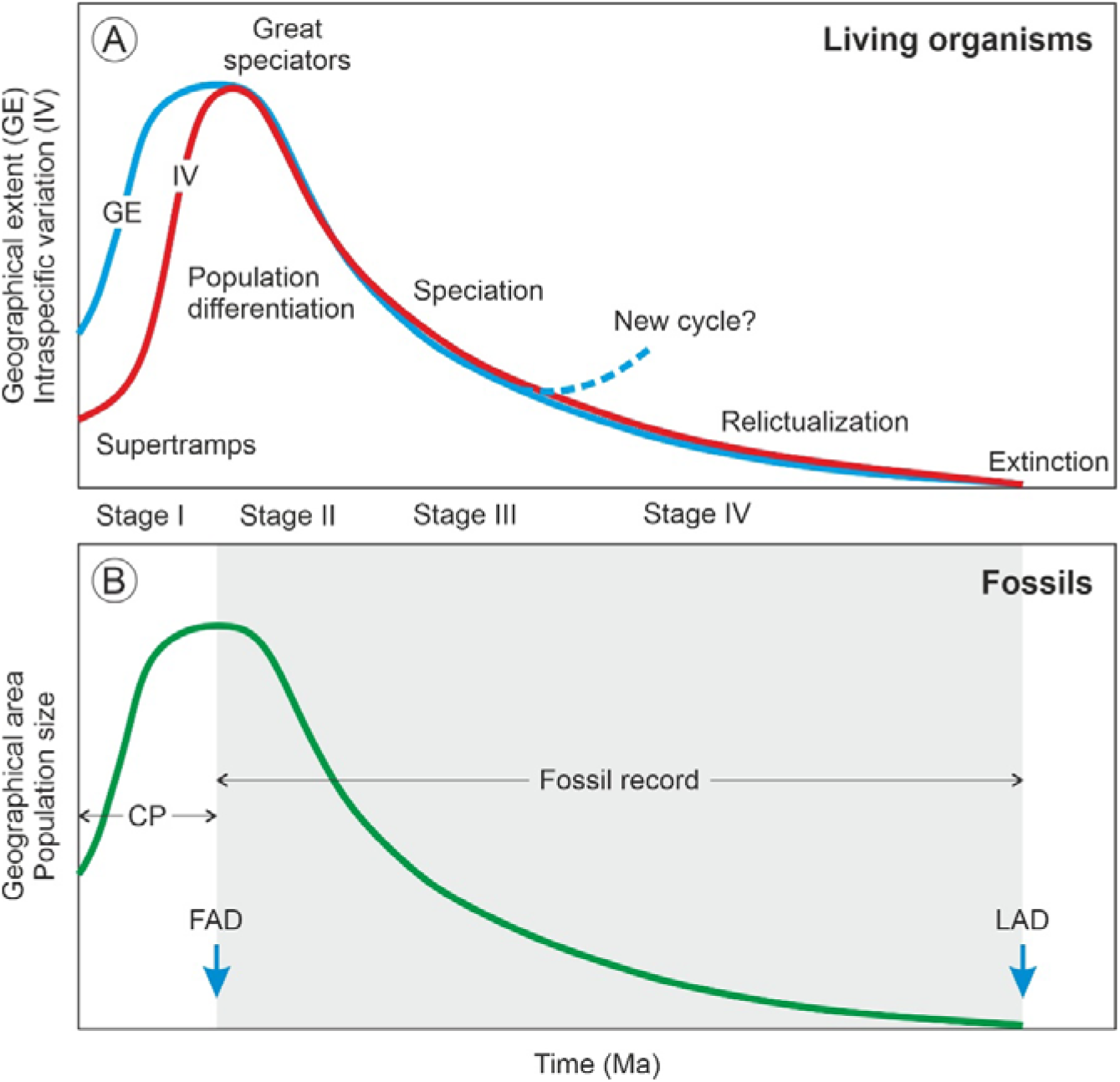
The taxon cycle and its fossil expression. A) Variations in geographical extent and intraspecific differentiation along the stages of the taxon cycle (redrawn from Pepke et al., 2019). B) Variations in the geographical area of fossils according to the asymmetric model (Lopez-Martinez, 2009). The stages represented in the fossil record (gray area) and the condensed phase (CP), which is apparently unrecorded, are indicated. FAD, first appearance datum; FAD, last appearance datum.

Originally, taxon cycles were defined to explain the biogeographical patterns of island biotas (Wilson, 1959), and although Ricklefs & Bermingham (2002) pointed out that they may also occur in continental environments, archipelagos remain the most common targets. Special emphasis has been placed on animal groups (crustaceans, insects, reptiles, birds) from tropical islands and archipelagos (Caribbean Antilles, Indonesia, Melanesia, New Guinea, Philippines, Madagascar) (e.g., MacLean & Holt, 1979; Losos, 1992; Jones et al., 2001; Cook et al., 2008; Simberloff & Collins, 2010; Economo & Sarnat, 2012; Jønsson et al., 2014; Economo et al., 2015; Fuchs et al., 2016; Dalsgaard et al., 2018; Matos-Maraví et al., 2018; Oliver et al., 2018; O’Connell et al., 2019; Cozzarolo et al., 2019; Liu et al., 2020: Cognato et al., 2021). Few case studies are available from continental areas. For example, a meta-analysis of a widely distributed bird group (Campefagidae) from tropical Asia, Africa and Australia, including insular and continental areas, has been considered to be consistent with the taxon cycle concept (Pepke et al., 2019). Another case is the diversification of Central American salamanders of the supergenus *Bolitoglossa* (Plethodontidae) (Garcia-Paris et al., 2000). In plants, similar case studies seem to be lacking. In agreement with Sheh et al. (2020), the author has been unable to find examples providing empirical support for the predictions of the taxon cycle theory. A single molecular phylogenetic study on the plant tribe Gaultherieae (Ericaceae) has been found, suggesting that fruit color might follow a process similar to a taxon cycle (Lu et al., 2019).

Initially, the occurrence of taxon cycles was attributed to changing biotic interactions, notably competition and predation, rather than to shifts in environmental drivers (Wilson, 1961; Ricklefs & Cox, 1972). Later, it was realized that the estimated duration of taxon cycles in Caribbean birds, based on molecular phylogenetic analyses, was on the order of 10^5^-10^7^ years (Ricklefs & Bermingham, 2002), which is much longer than the period of most climatic drivers of cyclic nature, especially the Pleistocene glacial-interglacial cycles, which have periods of 0.02-1-0.1 million years (my) (Hays et al., 1976). Therefore, the idea of a biotic origin and control was reinforced. Further studies using similar methods estimated taxon-cycle periodicities of ∼5 my for Indo-Pacific birds (Jønsson et al., 2014; Pepke et al., 2018) and reaffirmed the idea that Pleistocene climatic cycles could have not been important drivers. However, these authors suggested that other environmental shifts of greater periodicities, such as plate collision or orogenesis, could have been involved (Pepke et al., 2019). It has also been suggested that biotic drivers play a major role in the expansion phase, whereas abiotic drivers are more influential in the retraction phase (Žliobaitė & Stenseth, 2017). In all these works, the duration of taxon cycles was deduced from phylogenetic divergence times, which are usually estimated using molecular clock assumptions or are modeled using a variety of indirect methods (Ho, 2020). Therefore, according to Parenti & Ebach (2013), phylogenetic divergence times are hypotheses, not empirical evidence.

The fossil record could provide straightforward evidence and more reliable chronologies, but unequivocal fossil evidence for taxon cycles is still lacking. Range expansion-contraction cycles have been observed in the fossil record (Foote, 2007; Foote et al., 2007; Liow & Stenseth, 2007; Žliobaitė & Stenseth, 2017) but have not been analyzed from a taxon-cycle perspective (Pepke et al., 2019). According to Lopez-Martinez (2009), this is due to the difficulty of detecting the chronological and geographical origin of a species. Some paleontologists argue that the waxing and waning of fossil species follow symmetric trends (Foote, 2007) but Lopez-Martinez (2009) considers the taxon-cycle model as a good example of a time-asymmetric biogeographical and evolutionary process, as the initial dispersion phase (stage I) is usually much faster – and, hence, much more difficult to document in the fossil record – than the ensuing contraction/diversification and further extinction phases (stages II to IV). As a result, fossils would be able to account mainly for phases II to IV, whereas phase I would remain hidden (likely condensed) and represented only by the seemingly synchronous appearance of the species in a more or less wide area (e.g., Brunet et al., 1995) (Fig. 1B). According to the punctuated equilibrium model of evolution, this asymmetry should be viewed as an intrinsic feature of the fossil record – and, therefore, of the evolutionary process itself – rather than an imperfection of the fossil, as usually considered (Eldredge & Gould, 1972; Gould & Eldredge, 1977). In summary, according to the asymmetric model, the last appearance datum (LAD) of a fossil reliably records its extinction but its first appearance datum (FAD) does not record its actual time and place of origin but of its initial spreading (Fig. 1B).

*Pelliciera* Planch. & Triana (Triana & Planchon) – also reported as *Pelliceria* in some early publications – is a genus of Neotropical mangrove trees of the family Tetrameristaceae – formerly in the Theaceae or the Pellicieraceae, as reported in a number of classical papers – which has traditionally been considered monotypic (*P. rhizophorae*) but has recently been split into two species: *P. rhizophorae* (Planch. & Triana) N.C. Duke and *P. benthamii* Planch. & Triana (Duke, 2020). These species are currently restricted to a small patch along the Caribbean and Pacific coasts of Central America and northwestern South America – called here the present *Pelliciera* range or PPR (Fig. 2) – which has been considered to be a relict of the larger, nearly pan-Neotropical distribution attained by *Pelliciera* during Tertiary times (Wijmstra, 1968; Graham, 1977, 1995; Rull, 1998, 2001).

**Figure 2.**
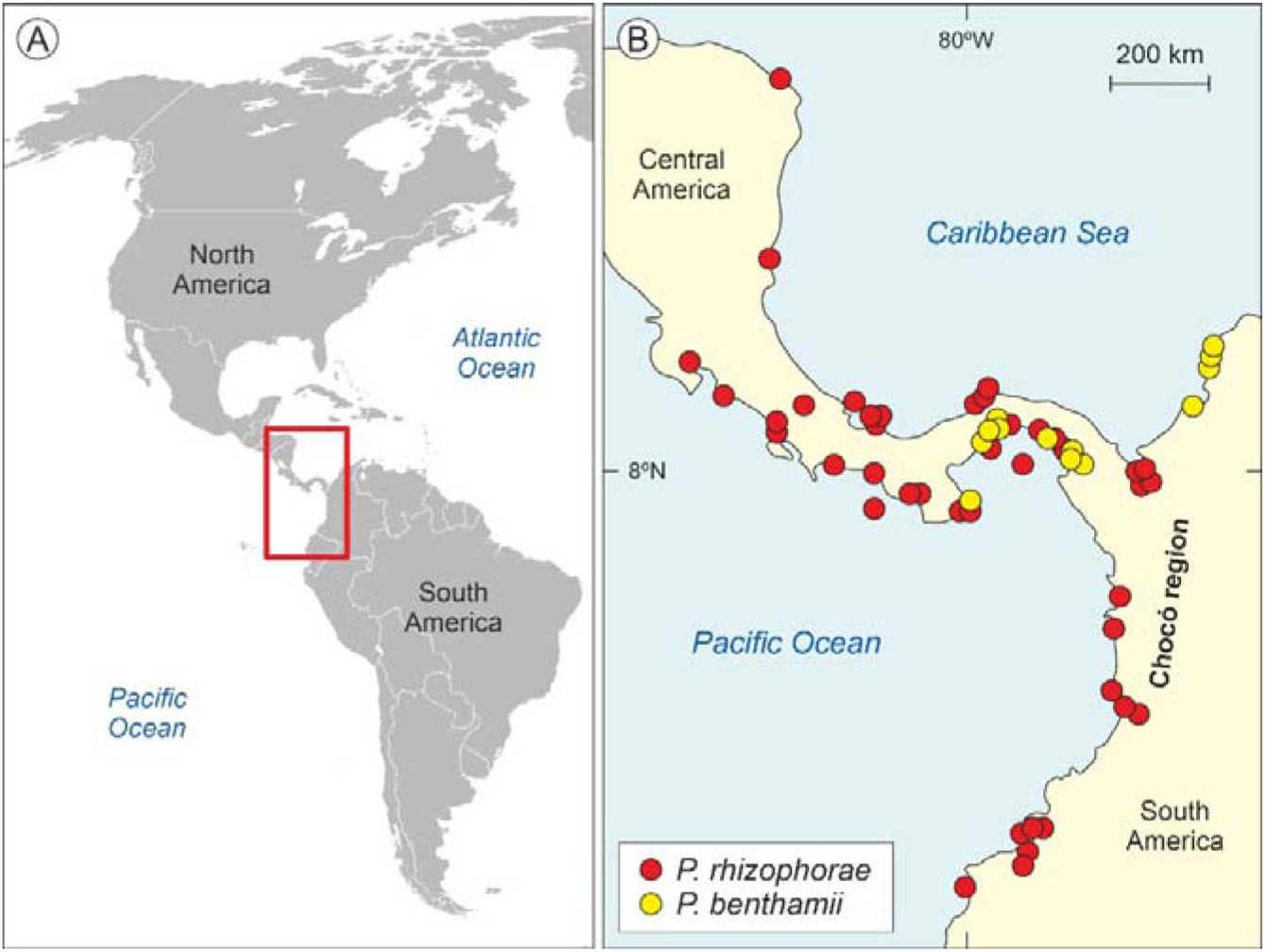
Present distribution of *Pelliciera* species. A) Map of the Americas with the distribution area of *Pelliciera* highlighted by a red box. B) Close up of the occurrence patterns of the two *Pelliciera* species (modified from Duke, 2020). The Colombian Chocó region is highlighted because it is one of the most humid regions of the world, with precipitation values up to 13,000 mm y^-1^ (Yepes et al., 2019).

Presently, *Pelliciera* is rare and restricted to sites with low or moderate salinity in the understory of *Rhizophora*-dominated mangrove forests (Dangremond et al., 2015). Recent autecological studies have shown that *Pelliciera* is highly sensitive to light intensity and salinity, and the combination of high levels of these environmental stressors leads to increased mortality, lower photosynthesis rates and reduced growth. When this species grows in shade conditions, however, it can tolerate high salinities, which suggests that light intensity is the main limiting factor. As a result, this taxon is unable to establish in sites with an open canopy and grows in the understory beneath the canopy of other tree species that, in the case of Central America, is provided by *Rhizophora mangle*, which is more tolerant to environmental stressors (Dangremond et al., 2015). It has also been reported that the scarcity of nutrients such as nitrogen and phosphorus can limit the growth of *Pelliciera*, leading to the development of dwarf forms (Dangremond & Feller, 2014). In general, *Pelliciera* is considered to be a stenotopic taxon bearing a relatively narrow and specialized niche.

Reconstruction of past biogeographical patterns and trends of *Pelliciera* are based on its fossil representative, the morphological pollen species *Lanagiopollis crassa* (van der Hammen & Wijmstra) Frederiksen, which also appears in the literature under the synonym *Psilatricolporites crassus* van der Hamen & Wijmstra (Wijmstra, 1968). This pollen originated locally in the Early Eocene but did not attain significant abundances until de Middle Eocene (Germeraad et al., 1968), when it became the dominant tree in the Neotropical mangrove communities (Rull, 2022a). By the time, *Rhizophora* (represented by the fossil pollen *Zonocostites ramonae* Germeraad, Hopping & Muller), currently the most abundant mangrove-forming tree in Neotropical coasts, was still absent in the region and arrived later, in the Mid-Late Eocene, likely by long-distance dispersal from the Indo-Pacific region, crossing the Atlantic through the Tethys seaway (Takayama et al., 2021). *Rhizophora* acquired its present dominant status during the Eocene/Oligocene Transition (hereafter EOT), which represented a global disruption characterized by a rapid cooling (which inaugurated the Cenozoic icehouse Earth’s state) and a sea-level fall that heavily influenced the Earth’s biota in the form of enhanced extinction and intense biotic turnover (Coxall & Pearson, 2007; Hutchinson et al., 2021).

In the Neotropics, the EOT signified a revolution for mangrove communities as the former Eocene *Pelliciera*-dominated mangroves disappeared and this tree turned into a minor subordinate element of the *Rhizophora*-dominated communities. However, this did not lead to the extinction of *Pelliciera*. Rather, this taxon was much more widespread across the Neotropics after losing its dominance than it was before and is today (Graham, 1977, 1995; Rull, 1998, 2001), which represents a major biogeographical challenge. Several explanations have been proposed to account for the Miocene-present *Pelliciera* reduction, including climatic and/or salinity stress, sea-level shifts or competition with *Rhizophora*, among others (Wijmstra, 1968; Fuchs, 1970; Graham, 1977, 1995; Jiménez, 1984; Rull, 1998, 2001). However, there are some methodological weaknesses that should be addressed before analyzing the potential causes for the *Pelliciera* biogeographical trends. First, although the idea of a post-Miocene range reduction was based on the analysis of barely a dozen of fossil records, this view has perpetuated until today with no further reconsideration based on an updated fossil database. Second, this reduction is only part of the story about the Cenozoic range shifts of *Pelliciera*, as pre-Miocene evidence has also not been analyzed under the same premises of a representative spatiotemporal fossil record. Third, most pollen records on which previous hypotheses were based consisted of qualitative (presence/absence) and pseudoquantitative (abundant, common, scarce) records, and it has been demonstrated that quantitative data are essential to properly record and understand the evolution of Neotropical mangroves (Rull, 2022a).

This paper uses an updated fossil pollen database of almost 80 widely distributed qualitative and quantitative Neotropical pollen records to reconstruct the biogeographical trends of *Pelliciera* since its Early Eocene origin to the present. This analysis revealed the occurrence of a long-term expansion-contraction loop that would compatible with the concept of taxon cycle, sensu Wilson (1961). Therefore, this would be not only the first evidence for a taxon cycle in plants but also a strong support for a taxon cycle, in general, as it is based on empirical, straightforward and chronologically accurate evidence, rather than on phylogenetic assumptions and modeling. The biogeographical cycle identified in this way is discussed in ecological and evolutionary terms using the known ecological traits of the taxa involved, primarily *Pelliciera* and *Rhizophora*, under the above-mentioned predictions of the conceptual model of Wilson (1961) and further updates. External (environmental) drivers potentially involved in the *Pelliciera* taxon cycle are also discussed. Considering the present features of *Pelliciera* populations, in comparison with its previous biogeographical history, the possible future developments are also evaluated.

## 2. Materials and methods

The data used in this paper are from the available literature; more information exists in the databases of oil companies but it is not publicly available. Efforts to make this information public, as is the case of Germeraad et al. (1968) and Lorente (1986), are worth to be made. After a comprehensive literature survey, almost 80 fossil mangrove pollen records were identified with data useful to reconstruct the biogeographical history of *Pelliciera* and *Rhizophora* in the Neotropics from the Eocene to the present. Almost half of these sites (45%) have quantitative data (percentages) and slightly more than a quarter (26%) have only presence/absence records; the rest have subjective measures such as abundant, common or rare. Ages are provided as geological epochs (Eocene, Oligocene, Miocene, Pliocene, Pleistocene and Holocene). In some cases, there is not enough chronological precision to resolve between them and longer ranges are indicated (e.g., Oligo-Miocene, Mio-Pliocene). The lack of site coordinates in many original references has prevented precise estimations of range sizes using statistical methods (Darroch & Saupe, 2018; Darroch et al., 2020); Carotenudo et al., 2020).

## 3. Results

The results obtained are displayed in Figure 3. Raw data (Table S1) and detailed figures for each time interval (Figs. S1 to S5) are provided as Supplementary Material. During the Eocene (17 Sites, including the two labeled as Eocene/Oligocene), *Pelliciera* was common or abundant in most localities, with maximum percentages up to 60%. The distribution area was restricted to NW South America (presently Colombia and Venezuela), with one site in eastern central America (Panama) and another on the Caribbean island of Jamaica. *Rhizophora* was present only in six localities, attaining some relevance (<10%) in only one site from Panama. In the Oligocene and Oligo-Miocene (18 Sites), a significant abundance decline is observed in *Pelliciera*, which falls to values below 5%, except in two localities from Venezuela. *Rhizophora* shows a reverse trend and is common or abundant in a significant number of sites, attaining values between 50 and 90% in four of them, situated in Guyana, Mexico, Venezuela and Puerto Rico. Regarding the distribution, *Pelliciera* shows a dramatic expansion in both latitudinal and longitudinal senses, ranging from Mexico and Puerto Rico to Brazil.

**Figure 3.**
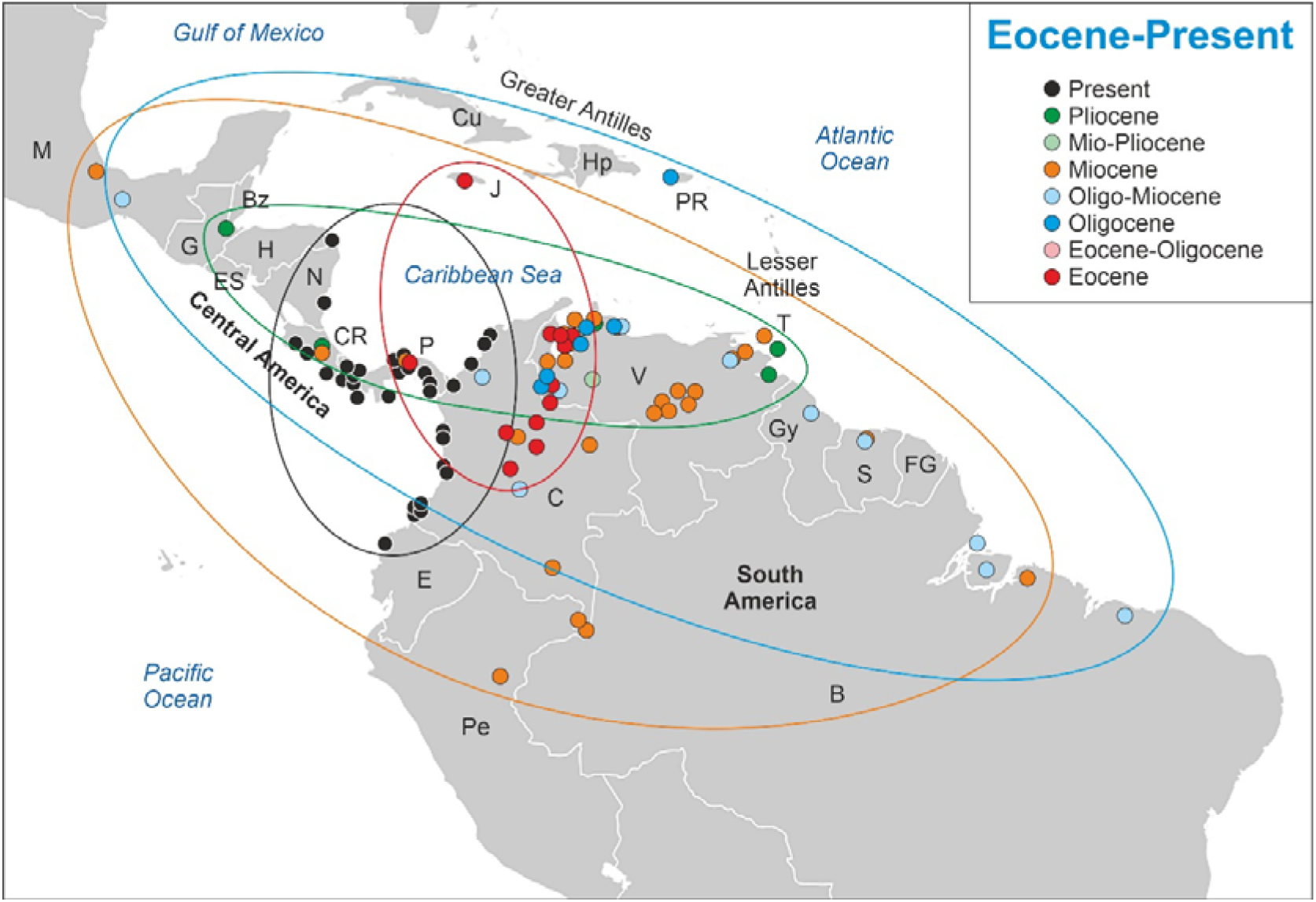
Graphical display of the results from Table S1 (Supplementary Material). Present *Pelliciera* localities have been taken from Fig. 2. Countries: B, Brasil; Bz, Belize; C, Colombia; CR, Costa Rica; Cu, E, Ecuador; ES, El Salvador; Cuba; FG, French Guiana; G, Guatemala; Gy, Guyana; H, Honduras; Hp, Hispaniola (Haiti and Santo Domingo); J, Jamacia; M, Mexico; N, Nicaragua; P, Panama; Pe, Peru; PR, Puerto Rico; S, Surinam; T, Trinidad & Tobago; V, Venezuela.

During the Miocene (36 sites), the decline in the abundance of *Pelliciera* continued and disappeared from 8 localities, reaching values of 3% only in five Venezuelan and one Panamanian site. The geographical distribution was similar to that in the Oligocene but with a slight displacement toward the SW. In contrast, *Rhizopohora* became abundant in most Miocene records with values up to 90%. In the Pliocene (8 localities, including some labeled as Mio-Pliocene), *Pelliciera* was only present and its range was restricted to the southern Caribbean margin (northern South America and Central America), whereas *Rhizophora* attained values of 70-100% in three sites and was present in others. No records exist for the Pleistocene and a few records are available for the Holocene (e.g., Horn, 1985; Jaramillo & Bayona, 2000), which are restricted to the PPR.

Shifts in the geographical range of *Pelliciera* represented in Fig. 3 can be subdivided into four main phases (Fig. 4). During the first phase (Eocene to Oligocene), the range expanded to most of the Neotropics but the populations experienced a significant reduction, which resulted in a more diluted distribution. This phase is called thinning expansion here, as the term dilution has already been used in biogeography for other processes (Keesing et al., 2006, 2010). The second phase (Oligocene to Miocene) was a displacement phase, where the range slightly migrated to the NW, with no significant differences with respect to the Oligocene situation. During the third phase (Miocene to Pliocene), the range underwent a major contraction toward the southern Caribbean margin. Finally, in the fourth phase (Pliocene to present) the range did not experience further reductions but showed a longitudinal contraction around Central America and a southward migration toward the Equator, which is considered a spatial reorganization. In summary, the range of *Pelliciera* significantly expanded from its original NW South American cradle until a Miocene maximum encompassing most of the Neotropics to initiate a retraction that ended in the present PPR, which is very similar in location and extent to the Eocene range. This biogeographical loop was accompanied by a dramatic decline in the *Pelliciera* populations, which was abrupt in the EOT and gradual between the Oligocene and the present (Fig. 5). If we consider that the Eocene *Pelliciera* mangroves attained their maximum extent in the Lutetian (Middle Eocene) (Rull, 1998, 2022a), the minimum duration of the overall *Pelliciera* biogeographical loop was approximately 45 my.

**Figure 4.**
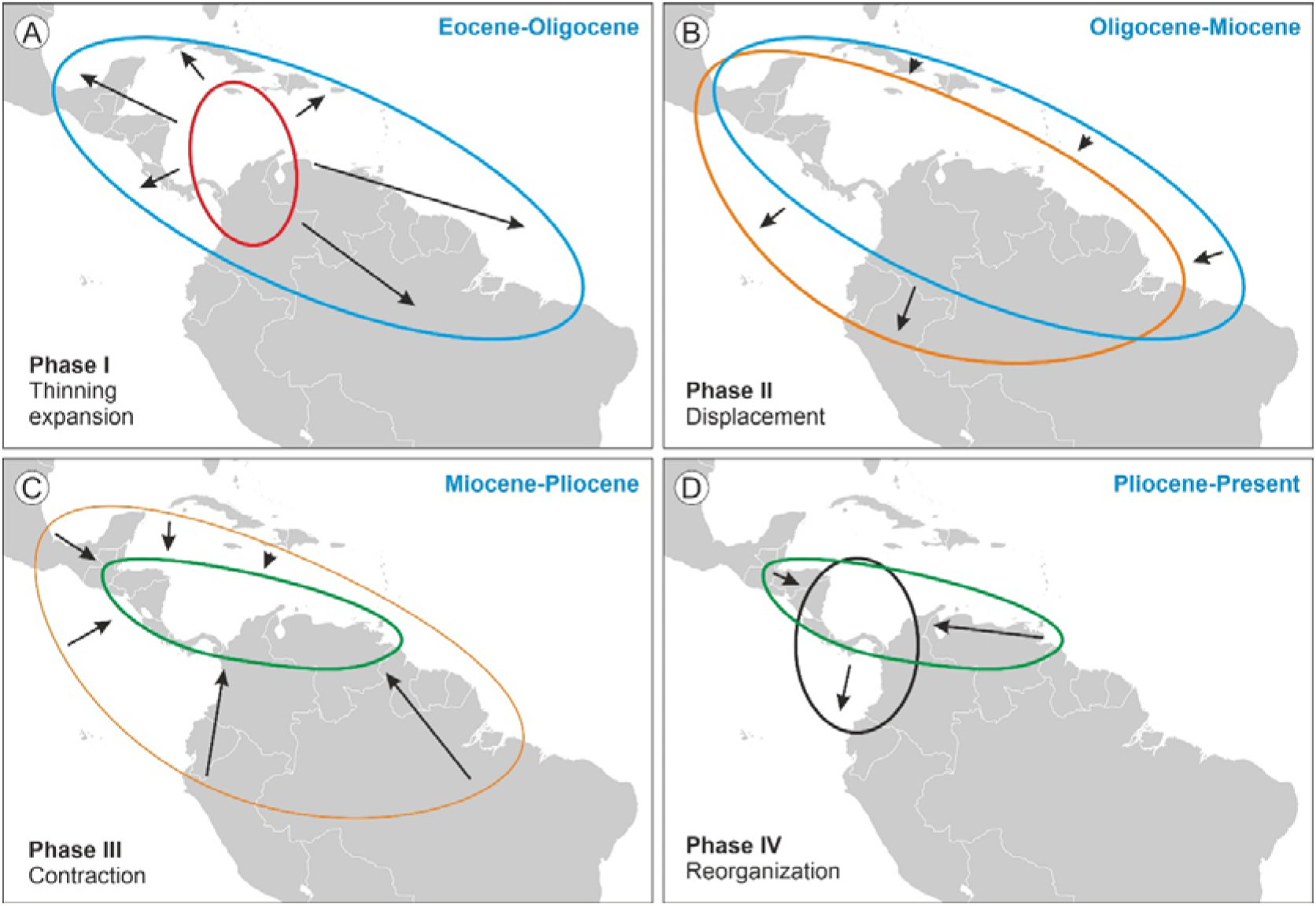
*Pelliciera* range shifts between the Eocene and the present, subdivided into four chronological phases.

**Figure 5.**
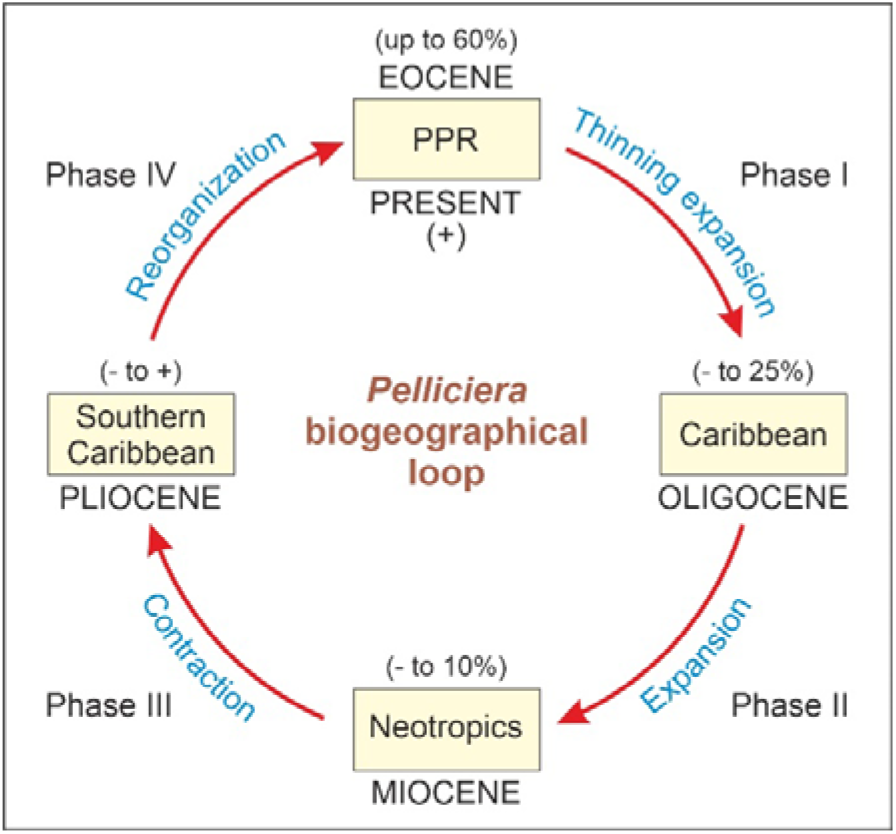
Diagram of the *Pelliciera* biographical loop with indication of the ranges of abundance of its fossil pollen in each geological epoch. PPR, present *Pelliciera* range.

## 4. Discussion

The *Pelliciera* loop, as documented in the fossil record, is straightforward empirical evidence for an expansion-contraction cycle, whose duration (∼4·10^7^ y) and biogeographical expression are consistent with the concept of taxon cycle, sensu Wilson (1961). Whether this cycle truly corresponds to a taxon cycle is discussed in more depth, with special consideration of the potential ecological and evolutionary implications.

First, it should be noted that the expansion phase (Eocene to Miocene) was much longer than expected under the asymmetric model, which postulates that phase I is too fast to be detectable in the fossil record (Fig. 1B). According to the asymmetric predictions, phase I should be condensed in the Eocene, when *Pelliciera* was already expanded across NW South America, Central America and the Greater Antilles (Fig. 3). To verify this possibility, the Eocene records (Fig. 2) have been subdivided into Early Eocene, Middle Eocene and Late Eocene (Table 1), revealing the occurrence of an intra-Eocene contraction-expansion loop (Fig. 6). According to this reconstruction, *Pelliciera* would have originated in the Early Eocene in western Venezuela and expanded relatively fast, attaining a maximum in the Middle Eocene, to further retreat (Late Eocene) to an area similar but not identical to its original range (Fig. 6). Therefore, the true stage I would have occurred between the Early and Middle Eocene. The exact duration of this loop is difficult to establish but a maximum estimate could be the total duration of the Eocene epoch, which is 22 my (∼2·10^7^ y). Using the same reasoning, stage I would have lasted a maximum of 10 my (10^7^ y).

**Figure 6.**
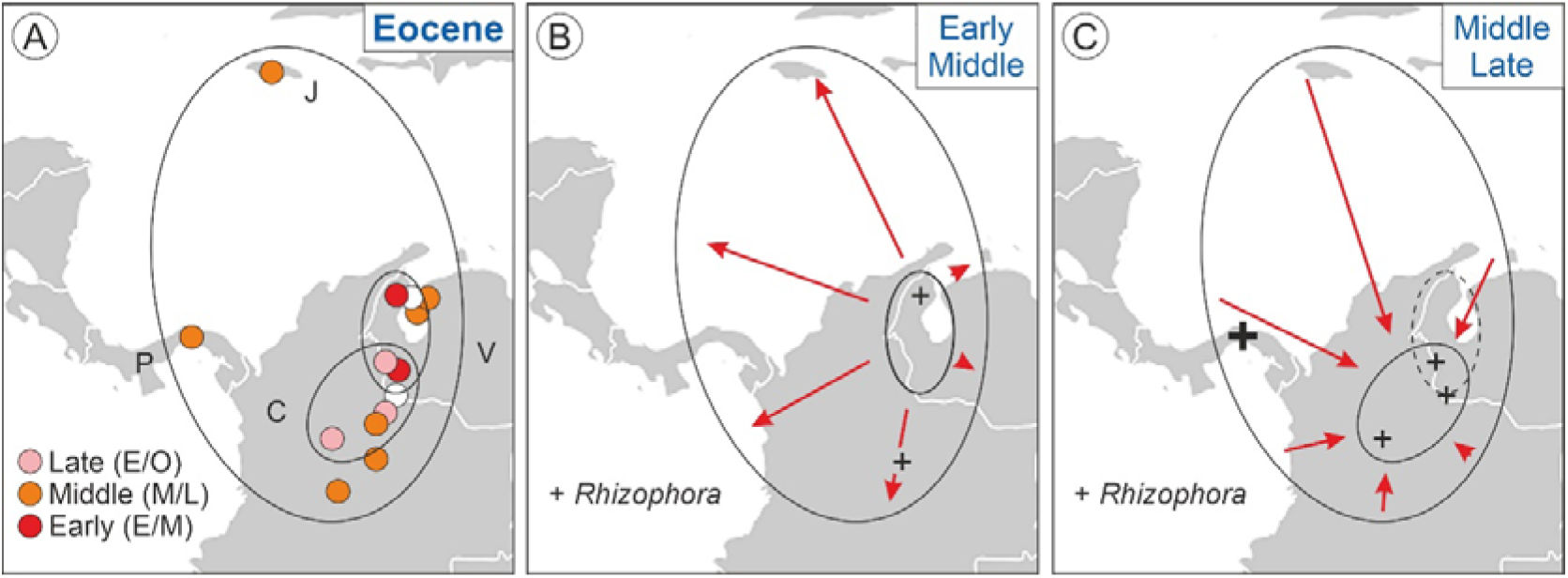
The Eocene expansion-contraction *Pelliciera* loop. A) Eocene localities (Fig. 3) subdivided into Early, Middle and Late Eocene ages (Table S1 of the Supplementary Material). Countries: C, Colombia; J, Jamaica; P, Panama; V, Venezuela. B) Early to Middle Eocene expansion. C) Middle to Late Eocene contraction. The initial Early Eocene range (A) is indicated by a broken line.

Evidence for population differentiation and eventual speciation should be sought on pollen morphology, which has demonstrated to be useful to differentiate extant *Pelliciera* species and subspecific taxa (Castillo-Cárdenas et al., 2014, 2015; Duke, 2020). The main diagnostic characters are size and exine (the outer pollen wall) sculpture. Fossil *Pelliciera* pollen (*L. crassa* or *P. crassus*) also exhibits significant morphological variability in these diagnostic characters from the Eocene to the Pliocene (Germeraad et al., 1968; Frederiksen, 1985; Lorente, 1986; Muller et al., 1987; Jaramillo & Dilcher, 2001). Unfortunately, no systematic records of this variability exist in the fossil record that enable to distinguish among possible taxonomic categories. This should be addressed in future studies, but with the available fossil evidence, the occurrence of different species and/or subspecies cannot be dismissed.

Regarding ecological preferences, most paleoecological studies using pollen rely on a reasonable degree of niche constancy over time (niche conservatism), especially at the genus level, in long-lasting communities (Wiens & Graham, 2005; Hadly et al., 2009; Wiens et al., 2010), which is the case of mangroves. Therefore, it is likely that fossil *Pelliciera* species were also stenotopic, as extant species are. This would be especially true in the Eocene, as the maximum recorded expansion of the *Pelliciera* range did not progress beyond tropical warm and wet climates, and is very similar to the present range, characterized by average temperatures of ∼27 °C and total annual precipitation values up to ∼3000 mm (Castillo-Cárdenas et al. 2015; Dangremond et al., 2015). It is especially noteworthy that extant *Pelliciera* grows around one of the most humid areas of the world, the Chocó region, (Fig. 2), with precipitation values reaching 13,000 mm y^-1^ (Yepes et al., 2019). During the Eocene, climates were significantly warmer than at present and global average temperatures were ∼8-14 °C above present temperatures (Westerhold et al., 2020), which would indicate macrothermal conditions for fossil *Pelliciera* species.

The first records of *Rhizophora* dated from the Middle Eocene and extended up to the Late Eocene in the form of scattered appearances, always around the initial and final Eocene ranges of *Pelliciera* (Table 1). The only exception is a Panamanian site (67), where *Rhizophora* reached 10% of the pollen assemblage during the Late Eocene. These occurrences represent the first stages of colonization of the Neotropics by *Rhizophora*, which coincides with the results of recent molecular phylogeographical studies that situate the origin of this genus in the Indo-Pacific region and its worldwide spreading in the Mid-Late Eocene (Takayama et al., 2021). According to the same authors, *Rhizophora* could have reached the Neotropics by the Atlantic Ocean, via the Tethys seaway. The use of this pathway cannot be supported or dismissed by paleogeographic reconstructions and fossil pollen records, as both Atlantic and Pacific seaways to the Caribbean were open for dispersal during the whole Eocene (Romito & Mann, 2020; Mann, 2021), and the fossil records do not show a clear Atlantic or Pacific pattern (Fig. 6). Once more, the only exception is site 67, which was situated in the Pacific island arc of the western Caribbean plate margin during the Eocene, which opens the door to an eventual Pacific dispersal pathway, but more studies are needed for a sound assessment.

The Oligocene to Miocene *Pelliciera* expansion (thinning expansion) started from a small area in NW South America, at the intersection between the Eocene and the present ranges (Figs. 4 and 6). Since the beginning, this expansion has followed the expansion of *Rhizophora*, the dominant mangrove tree. This change in dominance would have been greatly influenced by the EOT environmental disruption, in combination with biotic interactions. The EOT cooling would have affected a stenotypic macrothermic taxon such as *Pelliciera* and favored the development of the more eurytopic and climatically tolerant *Rhizophora*. In addition to its wider environmental tolerance, *Rhizphora* is an aggressive colonizer with a high dispersal potential, as its propagules are able to remain floating and viable for a year or more in saltwater. In contrast, the *Pelliciera* propagules have a maximum flotation period of barely a week and a maximum viability of roughly a couple of months, and their dispersal occurs primarily over short distances transported by coastal currents (Rabinowitz, 1978; Van de Stocken et al., 2019). The whole picture may suggest that the competitive superiority of *Rhizophora* could have led to the extinction of *Pelliciera*, but in contrast, this taxon not only survived but also expanded its range at the same pace of *Rhizophora* (Fig. 4). This could be explained by a combination of two types of biotic interactions known as facilitation and niche segregation.

Facilitation occurs when a species provides refuge to another in the face of environmental stress, predation or competition, thus allowing its survival (Boucher et al., 1982; Callaway, 1995; Stachowicz, 2001; Bruno et al., 2003). Niche segregation refers to the spatial, temporal or functional divergence of the niches of two competing species that allows the survival of both (MacArthur & Levins, 1967; Violle et al., 2011; Kosicki, 2022). As quoted above for extant mangroves, *Pelliciera* is highly sensitive to environmental stressors, and its growth is facilitated by *R. mangle*, which provides a favorable microhabitat in the mangrove understory (Dangremond et al., 2015). The maintenance of this specialized microhabitat could be explained by the fact that, although eurytopic generalists are apt to live in a wide range of environmental conditions – and, therefore, able to invade the specialist’s microhabitat – stenotopic specialists are more efficient within the restricted set of conditions in which they can develop (Futuyma & Moreno, 1988).

Using these ecological relationships as modern analogs for EOT mangroves, the *Pelliciera* to *Rhizophora* dominance shift without competitive exclusion could be viewed as a tradeoff in which the newly arrived eurytopic generalist (*Rhizophora*) outcompeted the resident stenotopic specialist (*Pelliciera*) in terms of dominance and, in exchange, facilitated a microhabitat to the looser, which “accepted” to play a secondary role in order to survive. In addition to survival, *Pelliciera* acquired the opportunity to expand its range across the whole Neotropics beyond its macroenvironmental niche boundaries, during the Oligocene and the Miocene. This would have been hardly possible without the facilitation of *Rhizophora*, as suggested by the permanent restriction of *Pelliciera* within its comfort zone (or the headquarters, in the words of Wilson, 1961) during the whole Eocene, when *Rhizophora* was absent. Given the dispersal mode of *Pelliciera*, typically over short distances through coastal currents, its expansion should have been by diffusion, that is, gradual migration across hospitable terrains, rather than by long-distance dispersal, which implies crossing inhospitable lands (Pielou, 1977). This suggests that *Rhizophora* must have been the pioneering colonizer, and once the mangrove community was developed and the microhabitat suitable for *Pelliciera* was created, this taxon would have been able to establish. Therefore, during its maximum Oligo-Miocene expansion, *Pelliciera* likely survived as a diffuse network of small populations restricted to favorable microhabitats, which is supported by mangrove fossil pollen records of this age, which are widespread but show very low pollen percentages, when present. This spatial arrangement and ecological dynamics fit with the concept of microrefugia, specifically the diffuse type, which promotes genetic differentiation among populations of the same species (Rull, 2009). Paraphrasing Fernández-Palacios et al. (2021), the ecological looser could have had unprecedented opportunities for diversification and become an evolutionary winner.

From an environmental point of view, the expansion of *Pelliciera* occurred in a phase of extended climatic stability spiked only by minor shifts (Westerhold et al., 2019), which suggests that biotic interactions were the main drivers in this part of the cycle. It is important to emphasize that the Oligo-Miocene spreading – whose duration is difficult to estimate due to the lack of enough temporal resolution in the fossil record – was the second expansion of *Pelliciera*, after the first occurred during the Early-Mid Eocene around the original range and its further Late Eocene contraction. Therefore, *Pelliciera* experienced at least two expansion/contraction cycles since the Eocene. The causes for the occurrence of the first cycle remain unknown but the second was likely triggered by the Late Eocene arrival of *Rhizophora*, which not only provided new spreading and diversification opportunities but also physically mediated the process, as a niche builder and an indirect dispersal agent (actually an ecological nurse). It is not possible to know whether *Pelliciera* would have become extinct without the arrival of *Rhizophora*, as its range was receding but, if so, *Rhizophora* could have provided the conditions for *Pelliciera* to overcome this bottleneck. Therefore, the ecological winner would have been not only the ecological nurse for the ecological looser but also its evolutionary rescuer. Considering the evolutionary dimension of these ecological interactions, *Rhizophora*, which likely began as a competitor, would have been much more than a lifesaver for *Pelliciera*, becoming with time a real sponsor/benefactor.

The Miocene-Pliocene contraction occurred after a significant cooling known as the Middle Miocene Cooling Transition (MMCT) (Westerhold et al., 2019). No evident changes were observed in the fossil pollen record, and hence, this contraction could have been more influenced by climatic than by biotic drivers. Obviously, the Pliocene restriction to the southern Caribbean margin was due to the local extinction of all Miocene *Pelliciera* populations outside this area, as predicted by the taxon cycle theory. It is possible that *Pelliciera* was unable to endure a second cooling, even under the protection of *Rhizophora*, and survived only in the sector of its range where temperature and precipitation remained favorable. This would be supported by the fact that the observed contraction had a clear latitudinal component. The reorganization that occurred between the Pliocene and the present is more enigmatic for two main reasons. On the one hand, the present distribution is based on actual records of living populations, while former distributions have been inferred from fossil pollen assemblages, which are subjected to a variety of taphonomic processes not affecting living plants. On the other hand, there is a gap between the Pliocene and the present, as no pollen records of *Pelliciera* are available for the Pleistocene and only a few Holocene records have been documented (e.g., Horn 1985; Jaramillo & Bayona, 2000). This prevent us from knowing how Pleistocene glaciations, the coolest phases of the whole Cenozoic era, affected the range of *Pelliciera*. Knowing the macrothermic nature of this taxon, it is expected that glaciations significantly affected its populations, but no evidence of this possibility is available to date.

An additional factor characteristic of the last millennia, which was absent in former geological epochs, is the presence of humans, whose influence on *Pelliciera* is largely unknown, except for the last decades. *P. rhizophorae* has been listed as “Vulnerable” – that is, under a high risk of extinction in the wild due to its small (500-2000 km^2^) and fragmented distribution area – in the IUCN Red List of Threatened Species (Polidoro et al., 2010; Blanco et al., 2012; Bhowmik et al., 2022). Urban expansion has been recognized as a major threat for *Pelliciera* populations, which are being heavily fragmented and threatened by habitat loss (Blanco-Libreros & Ramírez-Ruiz, 2021). From a biogeographical perspective, the available evidence suggests that, rather than shrinking the distribution area of *Pelliciera* as a whole, human activities have caused its severe fragmentation, which affects population viability and increases the sensitivity to extreme events and global warming (Blanco-Libreros & Ramírez-Ruiz, 2021). In the context of the taxon cycle, it could be asked whether the total extinction of *Pelliciera* is approaching and how human activities could contribute to accelerating this end. It is possible that, if the IUCN considers this view, *Pelliciera* would be transferred to the category Critically Endangered and considered a conservation priority. It is difficult to envisage the emergence of another evolutionary rescuer similar to *Rhizophora* that prevented the extinction of *Pelliciera* and promoted its expansion thus initiating an eventual third cycle. Humans could be able to do this through conservation/restoration actions, but we should seriously ask ourselves if we have the right to artificially preserve a taxon that is naturally headed to extinction (Rull, 2022b).

To summarize, the *Pelliciera* expansion-contraction cycles documented in the fossil pollen record since the Eocene have strong potential for being considered taxon cycles, sensu Wilson (1961), from chronological, biogeographical and ecological perspectives. Evolutionary predictions are more difficult to evaluate solely on the basis of the available pollen records and complementary evidence is needed for a sound assessment. Two main aspects remain to be analyzed in more detail in further studies, namely phase chronology and population diversification. Regarding the first, more precision is needed to accurately estimate the duration of the cycles and their respective phases. With reference to the second, more systematic pollen morphological studies, along with the use of molecular phylogenetic techniques, are required to document potential differentiation patterns among fossil populations. This paper has tried to maximize the utility of the information available in the literature in this sense. However, the existing studies were not aimed at demonstrating the existence of taxon cycles, which hinders more robust conclusions. With this in mind, further studies can be devised with the specific target of testing the evolutionary predictions of the taxon cycle model. Hopefully, this paper has provided a number of testable hypotheses that are worth exploring in this line. If confirmed, the *Pelliciera* cycles would be the first taxon cycles supported by straightforward empirical evidence and the first known taxon cycle for plants.

## Acknowledgments

No funding was received specifically for the development of this work.

